# Metagenomic contextualization of proteins with state space models

**DOI:** 10.64898/2026.07.07.736993

**Authors:** Nima Azbijari, Jacob H. Wynne, Kristopher Irizarry, Tanush Sistla, Andrew R. Thurber, Maude David

## Abstract

Genomic foundation models have emerged as powerful tools for learning representations of biological sequences, but existing approaches predominantly rely on Transformer architectures, whose quadratic attention complexity limits their ability to capture long-range genomic context. This limitation is particularly relevant for metagenomics, where sequencing data have increased more than 500-fold since 2011, generating increasingly large and diverse datasets. For example, gLM2, a Transformer-based mixed-modality genomic language model, was trained with a context window of 4,096 tokens, limiting its ability to capture extended genomic context within metagenomic contigs. Developing scalable architectures capable of processing longer sequences while integrating protein and nucleotide information is therefore essential for advancing genomic foundation models.

We introduce Nammu, a 167-million-parameter, bidirectional Mamba-based genomic foundation model that integrates protein and nucleotide sequences within a 20,000-token context window. Leveraging the linear scaling of state-space models, Nammu was trained using a curriculum learning strategy on the OpenMetaGenomic (OMG) corpus, first incorporating 64 million protein sequences followed by 32 million mixed-modality metagenomic contigs. Despite using approximately 37% fewer training tokens than gLM2, Nammu outperformed this Transformer-based model in taxonomic classification across all evaluated taxonomic ranks on a marine metagenomic dataset from the Critical Assessment of Metagenome Interpretation (CAMI). On the exact-feature-unmatched CAMI II Strain Madness test set, Nammu achieved higher all-contig genus-classification accuracy than Kraken2 (0.9182 versus 0.8142). Nammu further outperformed gLM2 (150M) in KEGG Orthology prediction using deep-sea metagenome-assembled genomes. Together, these results demonstrate the potential of state-space architectures for scalable, multimodal genomic representation learning and highlight their ability to support accurate taxonomic and functional predictions from metagenomic sequences.

## 1 Introduction

Rapid declines in sequencing costs have driven an unprecedented expansion of biological sequence datasets spanning a broad diversity of organisms (Consortium, 2015; Grigoriev et al., 2012; Mitchell et al., 2020). The increasing availability of these data has enabled the development of deep learning-based foundation models that learn representations directly from biological sequences, providing new approaches to understanding sequence structure and function. Protein language models, including ESM-2, ProGen, and ProteinBERT, have demonstrated their potential for protein structure prediction, functional annotation, and sequence generation (Lin et al., 2023; Nijkamp et al., 2023; Madani et al., 2020; Brandes et al., 2022). However, most protein language models process individual protein sequences independently, typically considering one open reading frame (ORF) at a time without incorporating its broader genomic context. Yet, protein function is often influenced by genomic organization, including gene neighborhoods, operon structure, and gene co-occurrence patterns (Mihelčić et al., 2019; Huynen et al., 2000; Pellegrini et al., 1999; Toibazar et al., 2024), suggesting that incorporating additional context could improve functional inference.

In parallel, nucleotide language models have demonstrated the ability to learn representations of genomic sequences, with applications ranging from regulatory element and variant effect prediction to genome-scale sequence modeling (Schiff et al., 2024; Nguyen et al., 2023, 2024; Brixi et al., 2026). Although these models can capture broader nucleotide sequence context, they do not explicitly integrate protein-level representations with their surrounding genomic sequences. Recent approaches have begun addressing this limitation by modeling proteins within their genomic context. For example, gLM represents genes within contigs using pooled protein language model embeddings to learn gene-gene relationships (Hwang et al., 2024), while gLM2 extends this approach by jointly modeling protein and nucleotide sequences, including coding and non-coding regions, strand information, and positional context (Cornman et al., 2024). These developments highlight the potential of integrating multiple sequence modalities to learn richer genomic representations.

Importantly, the genomic context surrounding protein-coding sequences also includes non-coding regions. Protein function is therefore influenced not only by its sequence and neighboring genes, but also by the surrounding regulatory landscape. Incorporating non-coding sequence context thus provides complementary information not captured by protein-centric models alone. Capturing this extended context requires models capable of processing sequences much longer than those of individual proteins. However, Transformer-based models (Vaswani et al., 2017), including gLM2, face computational limitations due to the quadratic scaling of attention with sequence length. For example, gLM2 was trained with a context window of 4,096 tokens, limiting the genomic context available within a single input. Alternative architectures, including Hyena-based models, state space models (SSMs), and hybrid approaches, have demonstrated the potential for more efficient long-range sequence modeling (Nguyen et al., 2023, 2024; Brixi et al., 2026; Sgarbossa et al., 2025; Wang et al., 2024).

Metagenomics is the culture-free sequencing of microbial community genomes, enabling the study of heterogeneous, uncultured microbial communities where a single sample can contain sequences from thousands of species (Wooley et al., 2010). From these shotgun sequencing approaches— initially dominated by short reads and now increasingly incorporating long reads—sequences are assembled into *contigs*, which reconstruct longer genomic fragments, enable recovery of genes, and provide the genomic context that can be leveraged to identify novel functions and link genes to organisms (Pavlopoulos et al., 2023). These contigs provide a rich source of sequence data well suited for unsupervised learning, as proteins are embedded within their native genomic context rather than treated in isolation. Assembled sequences can span tens of thousands of base pairs, capturing neighboring genes and regulatory elements. This context—such as synteny, gene fusion, and phylogenetic profiles—offers informative signals for protein function prediction beyond sequence similarity alone (Huynen et al., 2000; Pellegrini et al., 1999; Toibazar et al., 2024). These contextual signals provide a foundation for learning richer sequence representations in modeling frameworks, enabling more effective application to downstream biological tasks.

In this work, we present Nammu, a state space model (SSM)-based framework designed to overcome these limitations by extending the context window while jointly modeling protein sequences and surrounding non-coding DNA. We hypothesize that SSMs (Gu et al., 2022; Mehta et al., 2022; Gu et al., 2021; Gu and Dao, 2024) are well suited to capture the broader genomic context in which proteins are embedded. Specifically, Nammu builds on Mamba (Gu and Dao, 2024), an architecture with linear-time complexity *O*(*L*) that enables efficient processing of long metagenomic sequences. Previous SSM-based biological models, including ProtMamba, LC-PLM, and Caduceus, have demonstrated the potential of these architectures for modeling long protein or nucleotide sequences (Sgarbossa et al., 2025; Wang et al., 2024; Schiff et al., 2024). Building on these advances, we incorporate bidirectional modeling (Schiff et al., 2024) to capture both upstream and downstream dependencies between genomic regions. We train Nammu de novo using a two-stage pre-training strategy, beginning with protein sequences and subsequently transitioning to mixed-modality metagenomic sequences (Cornman et al., 2024; Wang et al., 2024). Together, these architectural and training strategies enable Nammu to learn representations of mixed-modality metagenomic contigs across substantially longer genomic contexts than are typically feasible with Transformer-based approaches.

## 2 Methods

### 2.1 Nammu Architecture and Training

#### Training Dataset: OpenMetaGenomic corpus

The model was trained using the Open-MetaGenomic (OMG) corpus as the primary data source (Cornman et al., 2024). We first initiated pre-training phase on protein sequences only, we use OMG_prot which is the protein-only subset of from the OMG dataset (phase 1 Figure 1). This subset of proteins created by the authors consists of 207M total proteins and was constructed by clustering sequences at 50% identity and removing singleton clusters (Cornman et al., 2024). For contig-level training we use the full OMG corpus which in total consists of 271M contigs, spanning 3.3B coding sequences and 2.8B intergenic sequences (Cornman et al., 2024).

**Figure 1:**
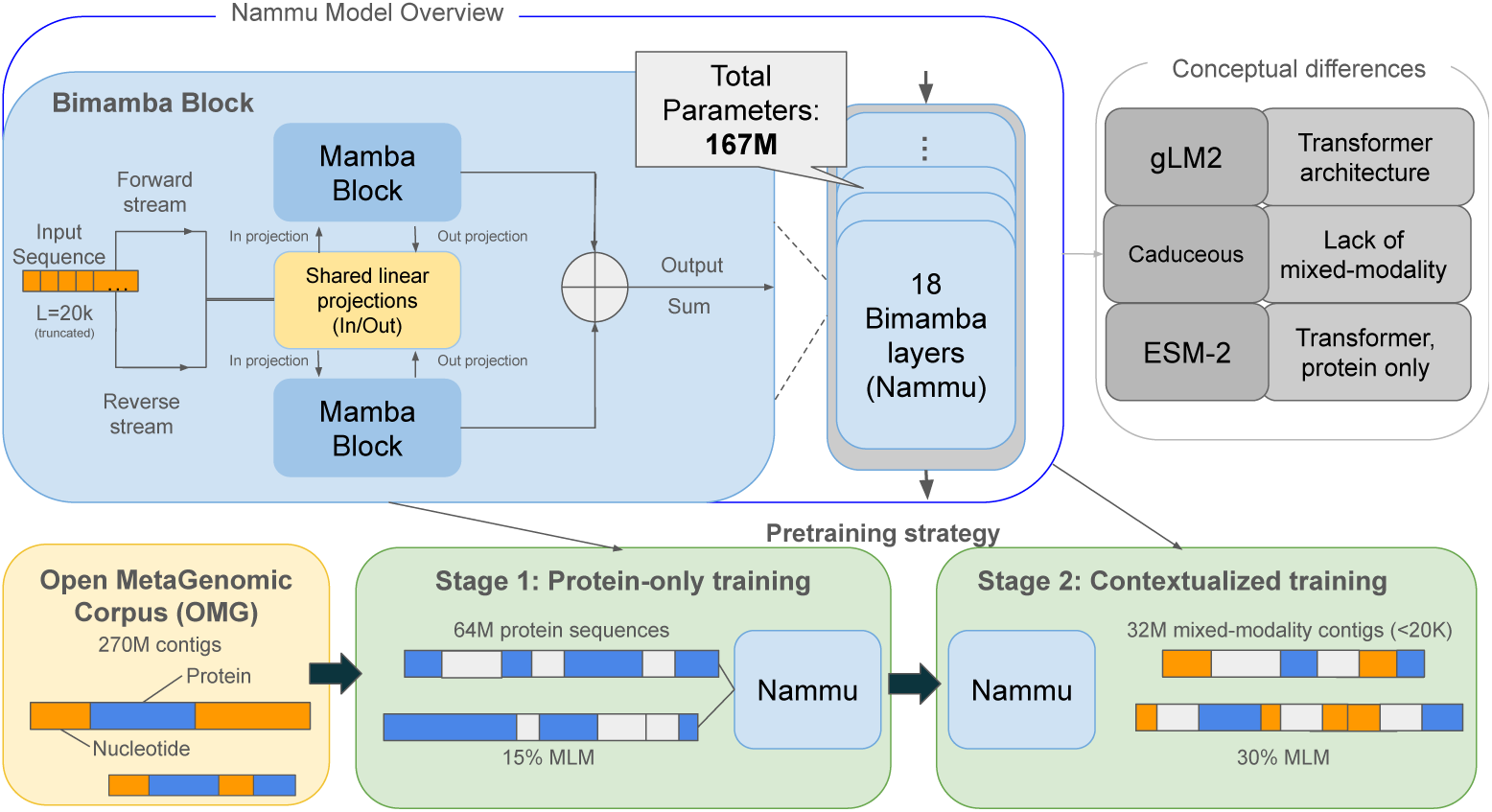
Overview of Nammu architecture and pre-training strategy. Nammu is a 167-million-parameter state space model comprising 18 bidirectional Mamba blocks with shared input and output projections. Pre-training was conducted in two stages, beginning with 64 million protein sequences (15% masking), followed by 32 million metagenomic contigs containing mixed protein and nucleotide sequences (30% masking).

#### Overview of the Nammu Architecture

The Nammu framework consists of 18 BiMamba blocks with shared linear projections between the forward and reverse streams (Figure 1). Each block processes the forward and reverse of the input and adds the results. Each BiMamba block also has an outer residual connection that adds the residual to the block output before being fed into the next block. The reversed input is only done across the sequence dimension. The SSM module in each Mamba block models each coordinate in an embedding with its own state space. This allows each coordinate along the sequence to be modeled as a signal in parallel with all other coordinates before having the learned signals mixed through the output projection.

#### Sequence Embedding Model and Training

We employed a curriculum learning strategy in which Nammu was first trained as a protein language model, then further pretrained on metagenomic contigs containing interleaved nucleotide and protein segments, extending prior staged bidirectional Mamba training approaches for protein sequence modeling (Wang et al., 2024). All models were trained with a fixed maximum context length of 20K tokens, with longer contigs truncated to this length. This context size was selected based on the contig length distribution of the OMG dataset, in which more than 95% of contigs are shorter than 20Kbp. Longer context lengths were associated with increased computational cost and training time with minimal increase in dataset coverage. Nammu was first pretrained on 64M protein sequences using a masked language modeling objective with 15% random masking. In the second phase of pre-training on mixed-modality sequences, we increased the masking rate to 30%, following the same strategy as gLM2.

Training was done in a self-supervised manner by masked language modeling (MLM) (Devlin et al., 2019). Given an input sequence consisting of proteins or DNA, along with strand orientation tokens as defined in Cornman et al. (2024), tokenization was performed at the resolution of the base pair, resulting in **x** = (*x*_1_*, x*_2_*, . . . , x_N_* ), where each **x**_i_ was either an amino acid, nucleotide, or strand orientation token. This is the same tokenization scheme as gLM2. A random subset *M ⊂ {*1*, . . . , N }* of token positions is masked. The model was trained to predict the original tokens at masked positions given the corrupted sequence **x̃**, where masked tokens were replaced by a special [MASK] token. The MLM loss was defined as:

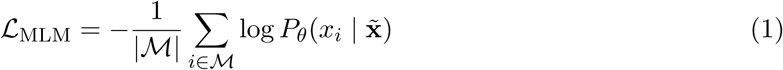

where *P_θ_*(*x_i_ |* **x̃**) was the probability assigned by the model with parameters *θ* to the true token *x_i_* at masked position *i*, conditioned on the full corrupted sequence **x̃**. These sequences were processed by stacks of bidirectional mamba blocks and trained with the masked language modeling objective as defined in Equation (1).

#### The BiMamba Block

Each BiMamba block in Nammu is based on Mamba-1 (Gu and Dao, 2024) and processes the projected input through a convolution and non-linearity before the SSM, alongside a parallel gating branch. The in and out projections are shared for both directions, while the SSM parameters remain independent. This weight-tying scheme reduces parameter count while preserving full bidirectional context. This bidirectional formulation was introduced in Caduceus (Schiff et al., 2024).

Protein pretraining was performed on 64M sequences from the 207M protein sequence OMG dataset (OMG_prot). Nammu was trained for 500K steps using AdamW with a batch size of 128 and a linear learning rate decay schedule. The model was then further pretrained on the OMG contig database for 250K steps using AdamW with a batch size of 128 and a cosine learning rate schedule ranging from 1 *×* 10*^−^*^3^ to 1 *×* 10*^−^*^4^ over 600K training steps, following gLM2. We stopped at 250K steps due to no improvement in the training loss. This stage used a masked language modeling objective with 30% random masking. All training was conducted on an NVIDIA GH200 system, with a total training time of approximately two months. Across both training stages, Nammu was trained on approximately 230B total tokens.

### 2.2 Multimodal Benchmarking: Taxonomic Classification Tasks

The Critical Assessment of Metagenome Interpretation (CAMI) provides community benchmarks based on metagenomic datasets with known source organisms and varying levels of biological complexity (Meyer et al., 2022). We leveraged 2 datasets from this Database to evaluate the potential of Nammu in performing taxonomic classification.

#### Taxonomic Benchmarking and Ablation Study Using the Marine Dataset of the CAMI II Dataset

First we used the Marine dataset from the CAMI II challenge, which consists of simulated metagenomic communities representing marine microbial environments. High-quality assemblies provided by CAMI were filtered to retain contigs *≥* 1,000 bp resulting in 101,968 contigs. Genes were predicted using Prodigal to extract nucleotide and protein sequences. CAMI ground-truth taxonomic identifiers were resolved using TaxonKit and matched to predicted genes and contigs. Sequences lacking informative species-level assignments were excluded, and the resulting dataset was characterized in terms of contig lengths, sequence retention, and taxonomic diversity across phyla and species. For this benchmark, we compared Nammu against gLM2, selected because it is currently the only other genomic language model designed to process mixed-modality sequences containing both protein and nucleotide information. For both models, training and test embeddings were extracted from intermediate layers (9th layer), and taxonomic classification was performed using a k-nearest neighbors classifier (k=10) with cosine similarity, implemented in scikit-learn (Pedregosa et al., 2011; Alain and Bengio, 2016). For contigs longer than its context window (4096 tokens), gLM2 classification was executed using a rolling window with a stride of 1024 positions. We then averaged the chunks together to aggregate the embeddings. We compared the performance of these approaches using Accuracy, Macro F1, Matthews Correlation Coefficient (Chicco and Jurman, 2020), Precision, and Recall. Performance metrics were calculated across taxonomic levels, including class, order, family, genus, and species to demonstrate performance as the task increases in granularity.

We also performed ablation experiments to assess the contribution of genomic context to taxonomic classification performance. Specifically, we independently shuffled the positions of entire coding sequences (CDS) or intergenic sequences (IGS) within each contig, preserving their internal sequences while disrupting their original genomic organization. Classification performance was then evaluated using the same approach as for the unmodified contigs, allowing us to assess the importance of coding and non-coding region organization for each model.

For this benchmark, we compared Nammu against gLM2, selected because it is currently the only other genomic language model designed to process mixed-modality sequences containing both protein and nucleotide information. We selected gLM2-150M as a comparator because its 150 million parameters are comparable to Nammu’s 167 million, allowing us to evaluate the two architectures at a similar model scale. For both models, training and test embeddings were extracted from intermediate layers (9th for Nammu, 15th layer for gLM2), and taxonomic classification was performed using a k-nearest neighbors classifier (k=10) with cosine similarity, implemented in scikit-learn (Pedregosa et al., 2011; Alain and Bengio, 2016). For contigs longer than its context window (4096 tokens), gLM2 classification was executed using a rolling window with a stride of 2048 positions. We then averaged the chunks together to aggregate the embeddings. We compared the performance of these approaches using Accuracy, Macro F1, Matthews Correlation Coefficient (Chicco and Jurman, 2020), Precision, and Recall. Performance metrics were calculated across taxonomic levels, including class, order, family, genus, and species to demonstrate performance as the task increases in granularity. We also performed ablation experiments to assess the contribution of genomic context to taxonomic classification performance. Specifically, we independently shuffled the positions of entire coding sequences (CDS) or intergenic sequences (IGS) within each contig, preserving their internal sequences while disrupting their original genomic organization. Classification performance was then evaluated using the same approach as for the unmodified contigs, allowing us to assess the importance of coding and non-coding region organization for each model.

#### Large-Scale Contig Classification and Comparison with Kraken2

In order to apply nammu in a more real life scenario, we launched an annotation at large scale using a second dataset form the CAMI II database, named strain madness, which deliberately contains many closely related genomes, making it useful for asking whether learned contig representations discriminate genera in a densely related, short-read-based assembly setting. We evaluated the frozen genus-labeled contig split from 100 samples: 11,878,846 training and 2,969,712 held-out test contigs, comprising 13 true genera designations, and 382 source-genome IDs in the test cohort. These are counts for our processed cohort, not the complete published CAMI II resource. Contigs were randomly partitioned between training and test (80/20). To examine possible overlap with pretraining data, we searched both partitions against the full OpenMetaGenomic (OMG) contig cache (Cornman et al., 2024) for exact matches to eligible complete predicted proteins (at least 100 amino acids) or intergenic segments (at least 150 nucleotides). Removing 3,198,104 training and 799,844 test contigs with at least one detected match, without reassigning contigs, yielded an *exact-feature-unmatched* sensitivity set of 8,680,742 training and 2,169,868 test contigs. We compared Nammu with Kraken2 (Wood et al., 2019) on the CAMI II Strain Madness dataset. Nammu embeddings were extracted from hidden state 9, mean-pooled, L2-normalized, and classified at the genus level using cosine nearest-neighbor search against training contigs (*k* = 1). Kraken2 was run using the 2025 Standard database, and taxonomic assignments were resolved to genus level. Both methods were evaluated on the same held-out contigs across 13 genera, with unresolved assignments counted as incorrect. Performance was assessed as a one-off genus-level annotation attempt using overall accuracy, with each contig weighted equally and unresolved assignments counted as incorrect.

#### Operon DB

Operons are groups of genes that are transcribed together, acting as an important component of bacterial gene regulation and function. To evaluate Nammu’s ability to distinguish intra-operonic from inter-operonic boundaries, we sourced experimentally verified operons from OperonDB obtained from 7 reference genomes (Pertea et al., 2009). For each intergenic region (IGS) flanked by two coding sequences (CDS1 and CDS2), we constructed an example consisting of the triplet (CDS1, IGS, CDS2) and a binary label indicating whether the two CDSs belong to the same operon. This resulted in 5,502 examples, comprising 3,636 intra-operonic examples and 1,866 operon boundary examples. The dataset was partitioned at the genome level into train, validation, and test splits to prevent leakage between splits. To evaluate Nammu’s performance on functional classification of operons, we extracted operons with well-annotated Enzyme Commission (EC) assignments from OperonDB. We assigned functional labels at the EC-2 (subclass) level, grouping operons by broad biochemical role, because operonic context is expected to influence function at a broad level. This yielded 1100 operons comprising 1564 CDSs from 22 genomes, spanning 15 EC-2 classes. The dataset was partitioned at the genome level into train, validation, and test splits.

#### Biosynthetic Gene Cluster Classification

A biosynthetic gene cluster is described as a set of genes clustered together on a segment of the genome that form a function together for producing metabolites that can be useful in medicine, algriculture, and other domains (Medema et al., 2015). We observe the distribution of sequence lengths in Figure **??** that these sequences can be long. Nucleotide only sequences are much longer and would require capturing more context while the mixed-modality sequences are compressed due to the translation for the coding regions. With the increased context window of 20K coupled with the sequence compression by representing coding regions as amino acids we can capture more context with Nammu. We utilize the MIBiG 4.0 dataset to measure BGC classification performance of Nammu (Zdouc et al., 2025). We filter the sequences to follow the OMG database filter criteria and only use sequences that have one associated BGC. This results in 1706 sequences spanned across 6 BGC classes from the MIBiG database.

#### Operon Boundary Classification

The ability to integrate protein-coding sequences (CDS) and intergenic sequences (IGS) within a longer genomic context offers opportunities to investigate relationships between neighboring genes and identify functional units, such as operons, that coordinate the expression of multiple genes. To evaluate this capability, we used experimentally verified operons from OperonDB to assess whether Nammu could identify operon boundaries, specifically the beginning and end of an operon (Pertea et al., 2009). We compared Nammu with gLM2-150M using two input configurations: CDS1+IGS (upstream coding context) and IGS+CDS2 (downstream coding context). These configurations were used to evaluate whether the sequence context surrounding an IGS could distinguish boundaries at the beginning or end of an operon from regions within an operon. Classification was performed using a k-nearest neighbors (kNN) classifier with cosine distance and distance-weighted voting. Performance was evaluated across 1,000 random train/validation/test splits (70/10/20), with the number of neighbors optimized on the validation set using MCC and final performance assessed on the held-out test set. Classification used a k-nearest-neighbour classifier with cosine distance and distance-weighted voting. We swept k over {1, 3, 5, 7, 11, 15, 21, 31, 51} and selected the value that maximized validation MCC. The selected classifier was refit on the combined train and validation set and evaluated on the held-out test set.

#### Functional Classification of Operons

Using the same OperonDB source, only operons with well-annotated EC assignments were kept, labelled at the EC-2 (subclass) level. Operons with empty labels were dropped and classes with fewer than 50 examples were removed. Embeddings were produced from the full operon context, with the hidden state pooled over the target CDS token positions. Embeddings were extracted once per model and reused across replicates. For each of 1000 replicates, examples were split 70/10/20 into train, validation, and test using class-stratified sampling on the EC-2 label. Classification followed the same kNN procedure and hyperparameter optimization strategy described above for operon boundary prediction.

#### Environmentally Sampled Deep-Sea Metagenomes

To evaluate Nammu’s performance on functional tasks, we tested its proficiency at identifying distinct KEGG Orthologs (KOs) using environmental sequencing, in contrast to CAMI benchmark datasets, which are simulated from reference genomes. For this task, we opted to use metagenome-assembled genomes (MAGs) from a highly complex, yet understudied environment: deep sea methane seeps (Han et al., 2023). Methane seeps represent unique microbial communities and function, given their chemical complexity. This results in a number of chemosynthetic and heterotrophic niches that remain poorly described with current methods. We used *>*3,000 MAGs generated by Han et al. (2023), spanning 16 methane seeps globally. From these MAGs, we predicted 6.2M open reading frames (ORFs) using Prodigal (Hyatt et al., 2010) and functionally annotated them using the tool DIAMOND (Buchfink et al., 2015) with the KEGG database (Kanehisa and Goto, 2000), using a cutoff of 90 percent sequence identity and an e-value lower than 1 *×* 10*^−^*^5^ to ensure highest confidence for KEGG ortholog predictions. This resulted in 67,172 ORFs that were functionally classified with a KO number, spanning 2,080 unique KOs.

## 3 Results

### 3.1 Nammu: Architecture and Training of a Long-Context Multimodal State Space Model

Biological sequence modeling has been dominated by transformer-based architectures (Munsamy et al., 2026; Hu et al., 2022; Hayes et al., 2025; Cornman et al., 2024; Lin et al., 2023), despite their quadratic scaling with sequence length. In contrast, state space models provide linear-time alternatives that enable modeling over substantially longer contexts (Sgarbossa et al., 2025; Schiff et al., 2024). Here, we introduce Nammu, a Mamba-1–based sequence model for mixed-modality metagenomic inputs (Gu and Dao, 2024). Nammu uses a BiMamba architecture (Schiff et al., 2024) consisting of 18 bidirectional Mamba blocks with shared projections across forward and reverse sequence directions, enabling efficient modeling of genomic contexts up to 20,000 tokens (Figure 1). The model was trained in two successive stages, beginning with 64 million protein sequences and subsequently extending to 32 million metagenomic contigs containing both protein-coding and intergenic regions. Across both stages, Nammu was trained on approximately 230 billion tokens. Across both pre-training stages, Nammu, comprising 167 million parameters, was trained on approximately 230 billion tokens.

### 3.2 Application of Long-Context Multimodal Models to Taxonomic Classification

#### Taxonomic Classification Using the CAMI II Marine Dataset

To evaluate Nammu’s ability to capture taxonomic information from metagenomic contigs, we first benchmarked its performance using the CAMI II Marine dataset. We retained contigs of at least 1 kbp with species-level taxonomic assignments, resulting in approximately 102,000 contigs spanning 246 species, 177 genera, and 37 classes. Open reading frames were predicted using Prodigal (Hyatt et al., 2010), and contigs were encoded as mixed-modality sequences containing amino acid representations of protein-coding regions and nucleotide representations of intergenic regions, following the OMG encoding scheme (Cornman et al., 2024). Using an 80/20 training and test split, we compared the taxonomic classification performance of Nammu and gLM2-150M across five taxonomic ranks: species, genus, family, order, and class (see Methods for details).

Nammu consistently outperformed gLM2-150M across all taxonomic ranks and evaluation metrics (Figure 2). The differences were particularly pronounced at broader taxonomic ranks. At the class level, Nammu achieved approximately 65% top-1 accuracy compared with 57% for gLM2, while the Matthews correlation coefficient (MCC) increased from approximately 0.28 to 0.43. Similar improvements were observed for macro F1, precision, and recall. Although classification performance decreased for both models at finer taxonomic resolutions, Nammu maintained higher performance across species, genus, family, and order classifications. These results indicate that Nammu’s pre-trained representations capture taxonomic information more effectively than those of gLM2 across multiple levels of taxonomic resolution. Nammu consistently outperformed gLM2-150M across all five taxonomic ranks and evaluation metrics (Figure 2). Improvements were statistically significant across all taxonomic ranks for macro F1, MCC, precision, and recall (Holm-corrected paired t-tests, *p <* 0.05, *p <* 0.01 or *p <* 0.001). For top-1 accuracy, significant improvements were observed at the genus, family, order, and class levels, while the difference at the species level was not statistically significant. The largest absolute improvements were observed at the class level, where Nammu achieved approximately 0.65 accuracy compared with 0.57 for gLM2-150M, and an MCC of approximately 0.43 compared with 0.28. At this taxonomic rank, Nammu also achieved substantially higher macro F1 (0.31 versus 0.16), precision (0.51 versus 0.31), and recall (0.25 versus 0.12). These results demonstrate that Nammu’s advantage extends beyond overall classification accuracy, with particularly pronounced improvements in class-balanced performance at broader taxonomic ranks.

**Figure 2:**
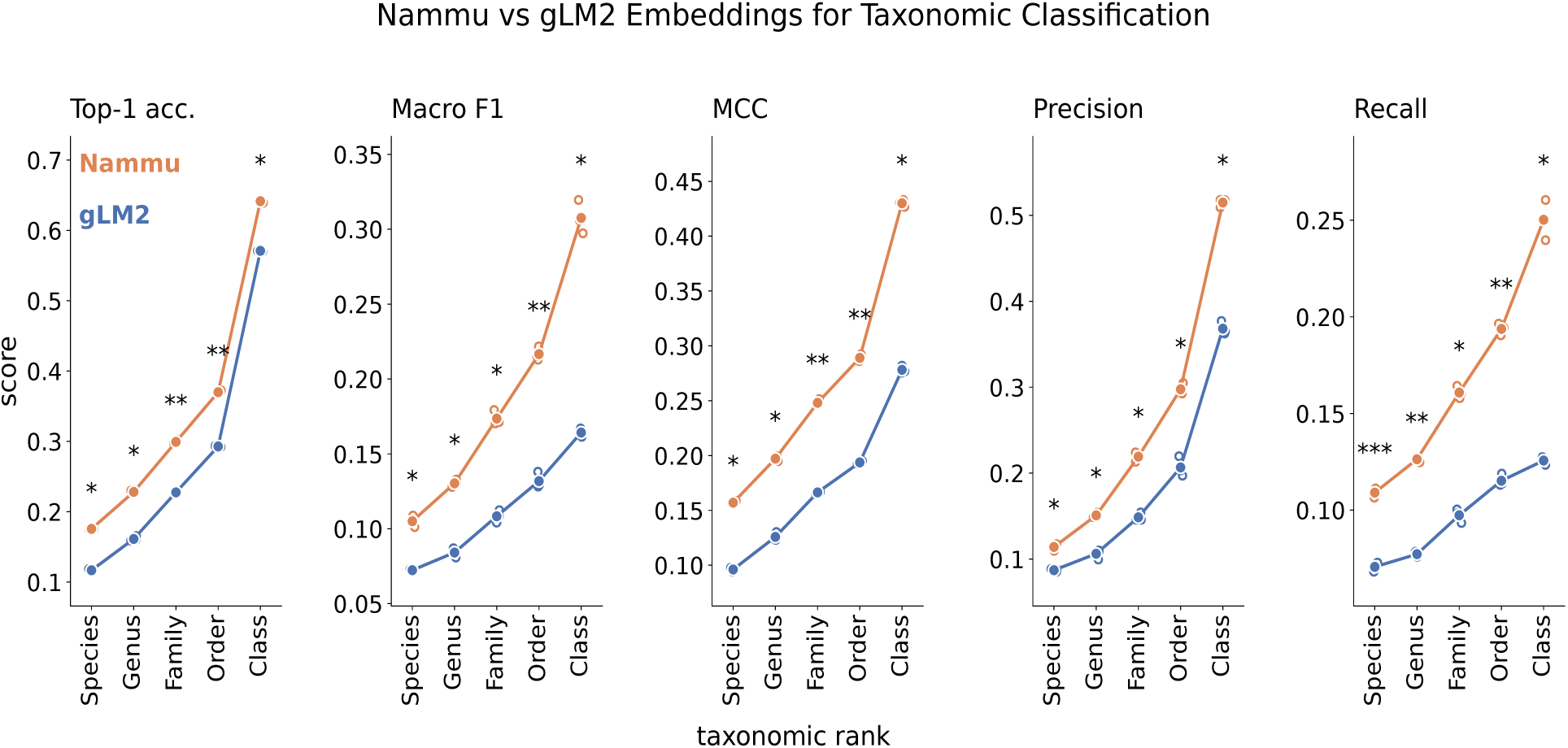
Taxonomic classification performance of Nammu and gLM2-150M on the CAMI II Marine dataset. Comparison of taxonomic classification performance across five taxonomic ranks (species, genus, family, order, and class) using embeddings generated by Nammu (orange) and gLM2-150M (blue). Performance was evaluated using five metrics: top-1 accuracy, macro-averaged F1 score, Matthews correlation coefficient (MCC), macro-averaged precision, and macro-averaged recall. Classification was performed using a k-nearest neighbors (kNN) classifier (k = 10) trained on pre-trained sequence embeddings, with an 80/20 training and test split. Results represent the mean across three random seeds, with error bars indicating standard deviations. Statistical significance between models was assessed using paired t-tests with Holm correction for multiple comparisons.

#### Taxonomic Classification and Ablations

We assessed gLM2 and Nammu embeddings of metagenomic sequences to predict taxonomy and multiple levels. Figure 2 illustrates embeddings from both models used to train a kNN classifier to predict taxonomy. Across the full set of sequences, we found that Nammu performs better for all taxonomic levels.

To take this further, we ran ablations to observe how different pieces of the sequence can impact classification of taxonomic class. Specifically, we tested the scenarios where either the IGS segments were shuffled by their position or the CDS segments were shuffled by their position. Figure 3 shows the results across each metric where the KNN is trained on non-shuffled sequences and the test sequences were shuffled. Interestingly, we only found that shuffling the IGS segments had any significant impact on performance but overall the shuffling seem to barely make an impact on classification.

**Figure 3:**
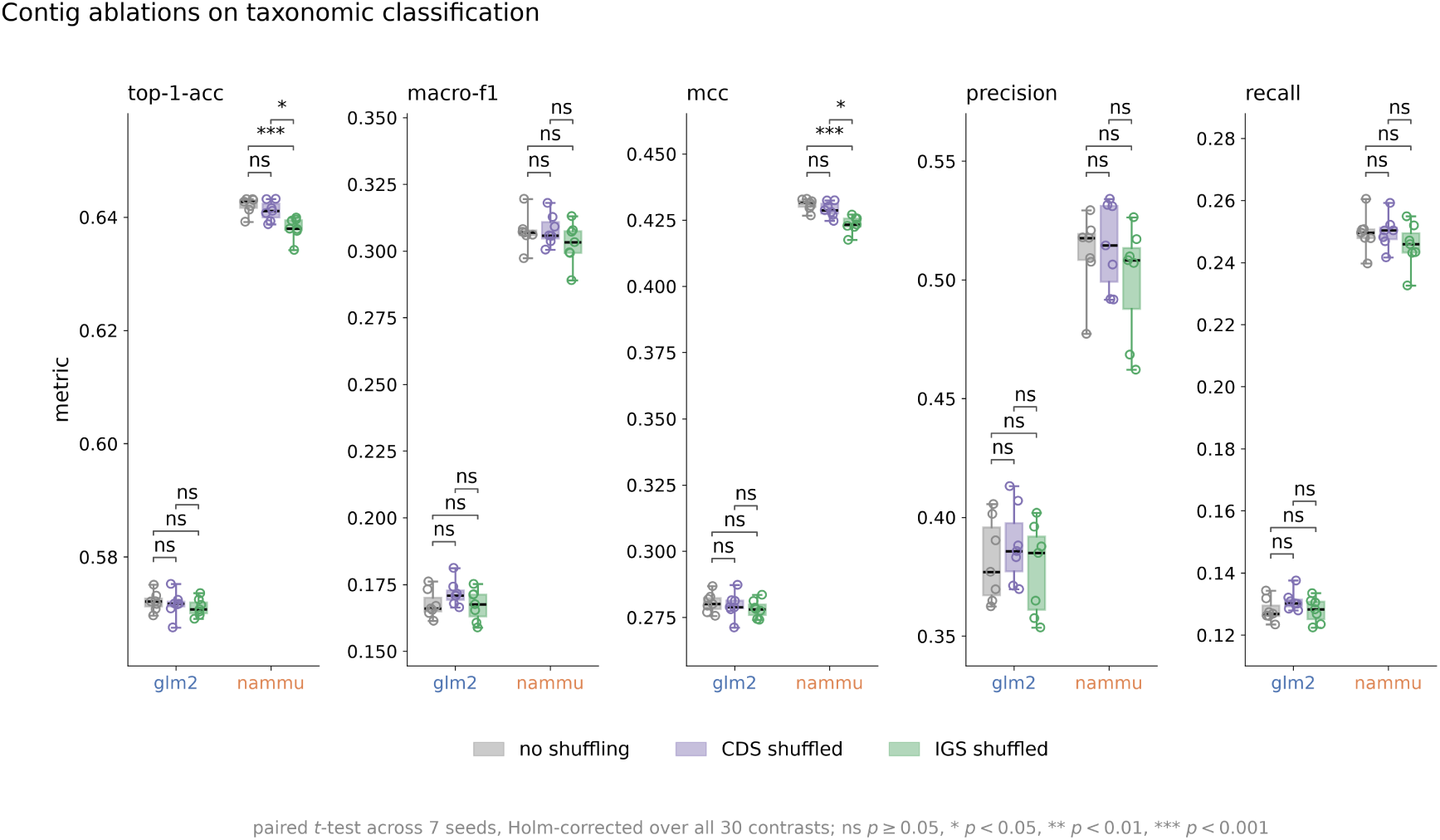
Performance on class level taxonomy for contigs of all lengths. Across 3 seeds per level, Nammu consistently performs better.

#### Large-Scale Contig Classification and Comparison with Kraken2

To evaluate Nammu’s performance on a larger and more challenging metagenomic dataset, we compared its genus-level taxonomic classification accuracy against Kraken2 using the CAMI II Strain Madness dataset. After excluding contigs with exact feature matches to the pre-training corpus, the test set contained 2,169,868 contigs representing 13 genera. Nammu achieved an accuracy of 91.82%, compared with 81.42% for Kraken2 /(a paired difference of +10.40 points) (Table **??**), corresponding to a 10.4 percentage-point improvement. These results demonstrate that Nammu’s learned representations can outperform the gold standard sequence-based taxonomic classification tool in this benchmark at the genus level.

#### Taxonomic Information Encoded in Nammu Representations

Given Nammu’s strong performance in taxonomic classification, we next investigated whether taxonomic information was reflected in its pre-trained embedding space by comparing the representations generated by Nammu and gLM2 using t-distributed stochastic neighbor embedding (t-SNE). We extracted embeddings from both pre-trained models without task-specific fine-tuning and visualized 101,968 metagenomic contigs, colored according to their class-level taxonomic assignments (Figure 2).

The two models exhibited distinct patterns of taxonomic organization. While gLM2 embeddings showed substantial overlap among taxonomic classes, Nammu embeddings displayed more pronounced grouping of several classes, including Gammaproteobacteria, Actinomycetes, and Flavobacteriia. Although considerable overlap remained, these observations suggest that Nammu’s pre-trained representations capture taxonomic structure more distinctly than those of gLM2 in this two-

Based on the results for class level taxonomic classification we also created a TSNE plots to observe clustering by class (Figure 4). We observe a clear clustering of taxonomy in Nammu’s embedding space while gLM2 produces mixed groupings.

**Figure 4:**
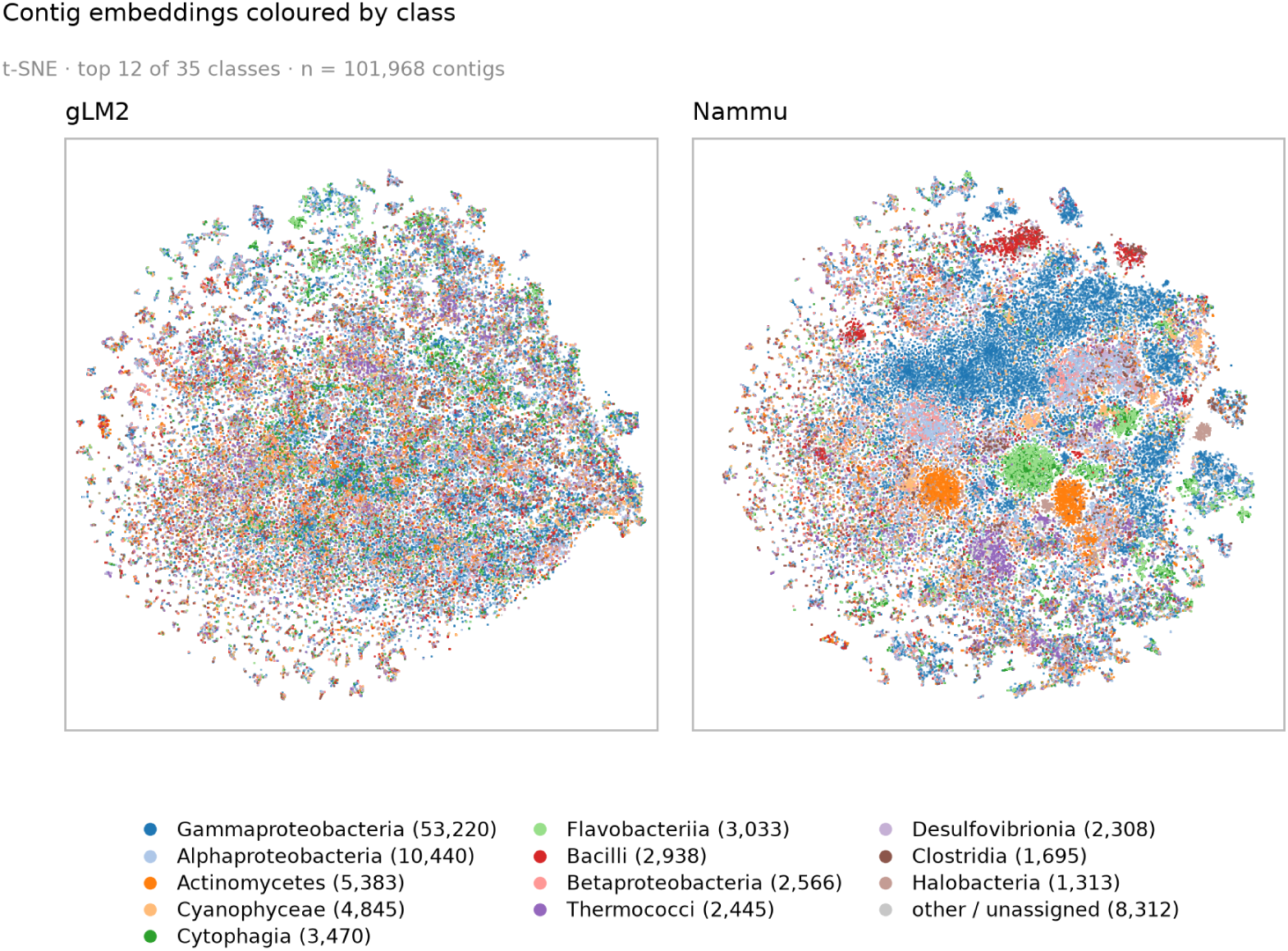
Embedding space for gLM2 embeddings and Nammu embeddings colored by class-level taxonomy. The embeddings from gLM2 are computed by a sliding window of size 4096 tokens with a stride of 2048. The windows are then averaged across token position before being averaged across the sequence dimension. Nammu uses no sliding window. Each point is a single vector representing the contig. Nammu exhibits better class grouping. dimensional visualization.

### 3.3 Application of Long-Context Multimodal Models to Functional Annotation

#### Biosynthetic Gene Cluster Functional Classification

Biosynthetic gene clusters (BGCs) consist of groups of neighboring genes that collectively encode pathways involved in the production of specialized metabolites, many of which have applications in medicine, agriculture, and biotechnology (Medema et al., 2015). As shown by the sequence length distributions (Supplement Figure 1), BGCs can span substantial genomic regions, making them particularly relevant for evaluating long-context sequence models. Representing BGCs as nucleotide-only sequences requires considerably longer input sequences, whereas mixed-modality encoding reduces sequence length by translating protein-coding regions into amino acids while retaining intergenic nucleotide sequences. This compression, combined with Nammu’s 20,000-token context window, allows larger portions of BGCs to be represented within a single model input, potentially capturing functional relationships among neighboring genes.

We evaluated Nammu’s performance in biosynthetic gene cluster (BGC) classification using the MIBiG 4.0 dataset (Zdouc et al., 2025). After applying the OMG database filtering criteria and retaining only sequences associated with a single BGC class, the final dataset comprised 1,706 sequences spanning six BGC classes.

Our objective is to predict to which of the 6 BGC classes from the MIBiG database the BCG belongs to. To do so, we embedded each BGCs from the MIBiG database, and used random forest, logistic regression, and k-nearest neighbors for the classification task. To process the sequences we use a sliding window (we defined the window to be 1/2 the context considered by each model, therefore Nammu uses its 20K window size with a stride of 10K. For gLM2-150M we used a window size of 4096 with a stride of 2048.) when embedding the contigs and we average over the windows to produce a single embedding to represent the sequence. In order to assess performance we made sure that taxa did not occur in both the train and the test set. Thus, if a particular taxa occurs in the training embeddings then that taxa will not be in the test embeddings. Additionally, we construct a k-mer baseline to benchmark both models (Supplement Table 1). Our results, shown in figure (Fig. 5, show that Nammu outperforms gLM2 in biosynthetic gene cluster classification in both nucleotide only and mixed-modality sequence representations.

**Figure 5:**
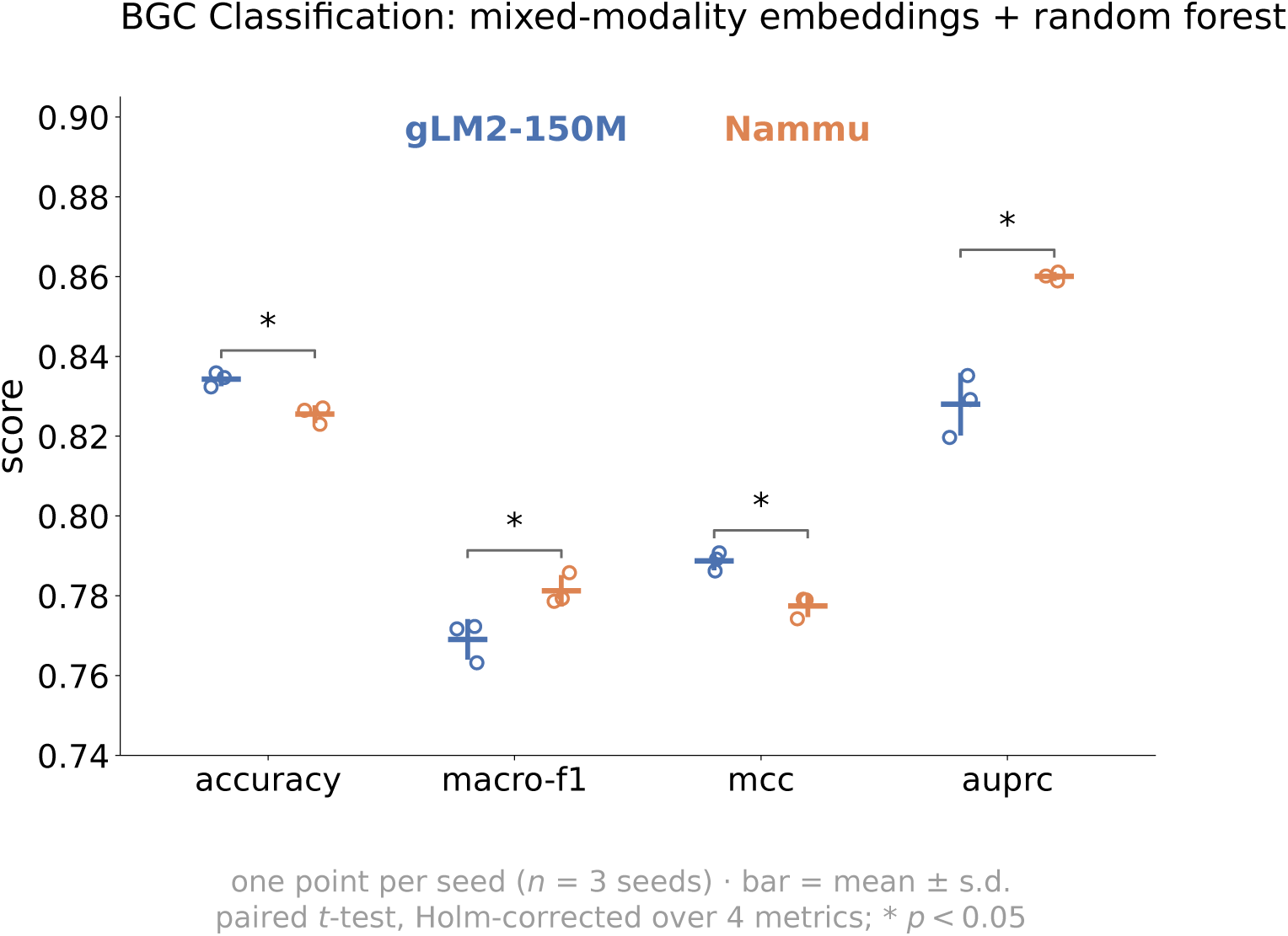
Comparison of Nammu and gLM2-150M for biosynthetic gene cluster (BGC) classification. Classification performance using mixed-modality embeddings extracted from layer 9 of Nammu and the middle layer of gLM2-150M, evaluated with a random forest classifier on the MIBiG 4.0 dataset (1,706 BGCs across six classes). Performance was assessed using accuracy, macro-F1, Matthews correlation coefficient (MCC), and macro-AUPRC. Statistical significance was assessed using paired t-tests with Holm correction for multiple comparisons. * indicates *p <* 0.05.

Using mixed-modality embeddings and a random forest classifier, Nammu achieved 82.6% accuracy, 0.781 macro-F1, 0.777 MCC, and 0.860 macro-AUPRC, compared with 83.4%, 0.769, 0.789, and 0.828, respectively, for gLM2-150M (Fig. 5). Although performance was comparable across most metrics, Nammu achieved a significantly higher macro-AUPRC (*p <* 0.05). These results highlight the value of mixed-modality representations for BGC classification and suggest that Nammu captures functional information that supports improved precision–recall performance across BGC classes.

#### Operon Boundary Prediction

For operon boundary classification, Nammu consistently outperformed gLM2-150M using both upstream (CDS1+IGS) and downstream (IGS+CDS2) sequence contexts (Table 2). Using IGS mean-pooled embeddings, Nammu achieved higher MCC for both upstream (0.648 vs. 0.599) and downstream (0.648 vs. 0.604) configurations, outperforming gLM2 in 91.8% and 86.8% of paired replicates, respectively. Improvements were also observed across all other evaluation metrics, including macro F1, precision, recall, and accuracy. In particular, Nammu achieved 83.8% accuracy for upstream and 83.7% for downstream context, compared with 80.7% and 81.2% for gLM2, respectively. These results suggest that Nammu’s representations of intergenic regions, informed by adjacent protein-coding sequences, are more effective at distinguishing boundaries within and between operons.

**Table 1:** Large-Scale Taxonomic Classification: Nammu versus Kraken2. Area under the precision–recall curve (AUPRC) for genus- and species-level taxonomic classification on the CAMI II Strain Madness test set.

| Method | Genus AUPRC | Species AUPRC |
| --- | --- | --- |
| Nammu | 0.88706 | 0.86500 |
| Kraken2 | 0.82991 | 0.60500 |

**Table 2:** Operon boundary classification using upstream and downstream coding sequence context. Comparison of Nammu and gLM2-150M using IGS-slice mean-pooled embeddings and kNN classification. Values represent mean (SD) across 1,000 replicates. Macro F1, precision, and recall are unweighted averages across classes. Bold indicates the best performance within each context.

| Model | MCC | F1 | Prec. | Recall | Acc. |
| --- | --- | --- | --- | --- | --- |
| *Upstream CDS + IGS* |  |  |  |  |  |
| Nammu | **0.648** (0.023) | **0.823** (0.012) | **0.817** (0.011) | **0.832** (0.012) | **0.838** (0.011) |
| gLM2-150M | 0.599 (0.025) | 0.794 (0.013) | 0.788 (0.012) | 0.812 (0.013) | 0.807 (0.012) |
| *IGS + Downstream CDS* |  |  |  |  |  |
| Nammu | **0.648** (0.024) | **0.822** (0.012) | **0.816** (0.012) | **0.832** (0.013) | **0.837** (0.011) |
| gLM2-150M | 0.604 (0.025) | 0.798 (0.012) | 0.791 (0.012) | 0.814 (0.013) | 0.812 (0.012) |

#### Enzyme Function Prediction Using Operon Context

To investigate whether Nammu captures functional information from the genomic context of operons, we evaluated its ability to predict enzyme functions at the second level of the Enzyme Commission (EC-2) hierarchy using experimentally annotated operons from OperonDB. Embeddings were extracted from target protein-coding sequences while incorporating their surrounding operon context. Nammu consistently outperformed gLM2-150M across all four evaluation metrics over 1,000 replicates (Table 3). Nammu achieved higher accuracy (0.860 vs. 0.713), balanced accuracy (0.847 vs. 0.690), macro F1 (0.848 vs. 0.693), and MCC (0.847 vs. 0.684). The largest absolute improvement was observed in MCC (+0.163), with Nammu outperforming gLM2 in every paired replicate across all four metrics. These results demonstrate that Nammu’s contextualized protein embeddings provide more informative representations for enzyme function prediction within operons.

**Table 3:** Full-operon functional prediction using Nammu and gLM2-150M embeddings with kNN classification. Values represent mean (SD) across 1,000 replicates. Bold indicates the best performance.

| Model | Accuracy | Balanced Acc. | Macro F1 | MCC |
| --- | --- | --- | --- | --- |
| Nammu | **0.860** (0.021) | **0.847** (0.024) | **0.848** (0.023) | **0.847** (0.023) |
| gLM2-150M | 0.713 (0.026) | 0.690 (0.030) | 0.693 (0.028) | 0.684 (0.029) |

#### Protein Embeddings for Functional Annotation of Environmental Deep-Sea Metagenomes

To evaluate Nammu’s ability to predict protein function in real-world sequenced environmental metagenomes, we benchmarked its performance on KEGG Ortholog (KO) classification using protein sequences extracted from deep-sea metagenomic ORFs. We compared Nammu with gLM2 using protein embeddings and a k-nearest neighbors (kNN) classifier, without incorporating surrounding genomic context. Across seven independent train-test splits, Nammu consistently outperformed gLM2 across all five evaluation metrics (Fig. 6). Nammu achieved higher top-1 accuracy (approximately 0.87 versus 0.86), precision (0.80 versus 0.78), recall (0.79 versus 0.77), macro F1 (0.78 versus 0.76), and MCC (0.87 versus 0.86). All improvements were statistically significant (Holm-corrected

**Figure 6:**
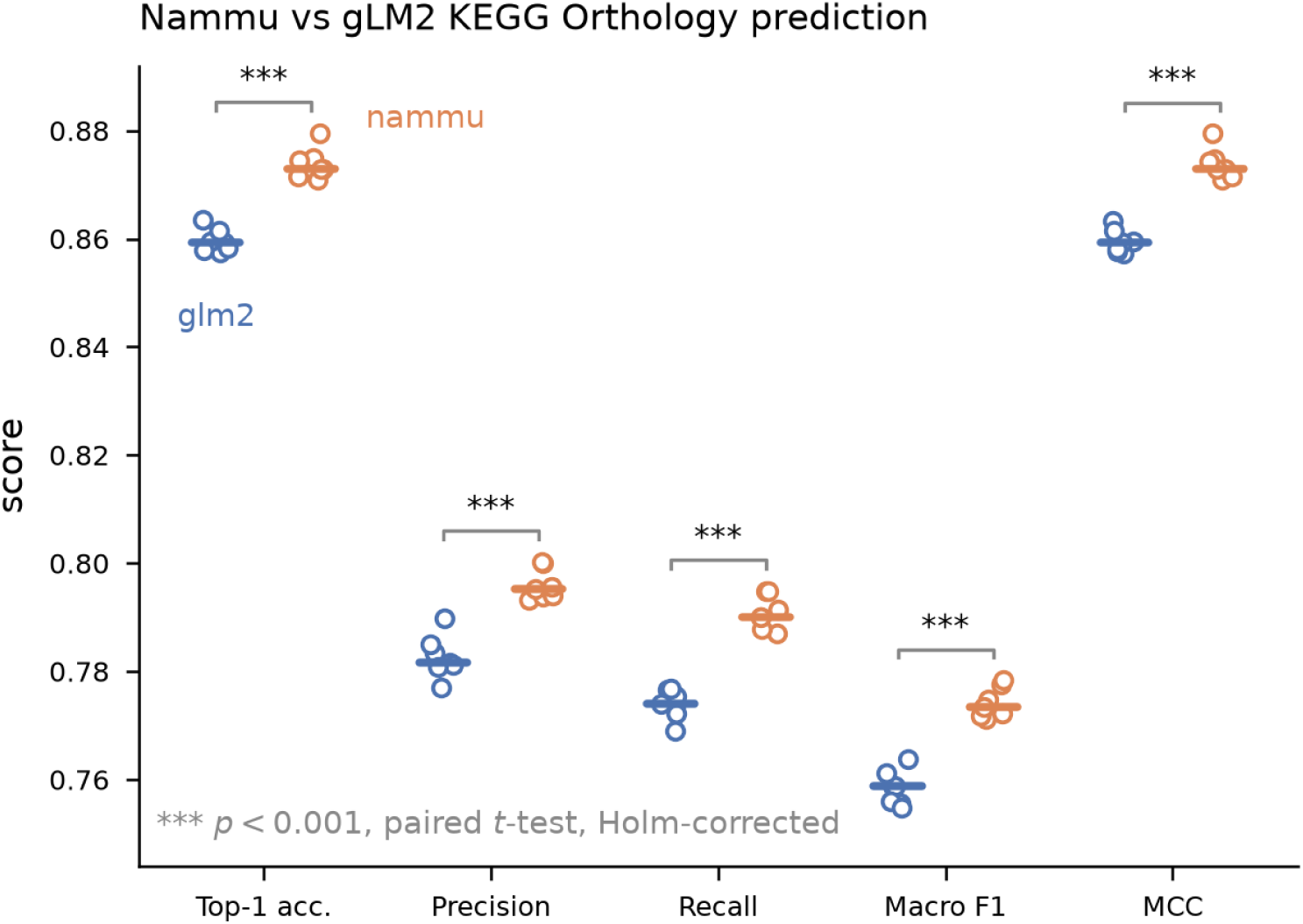
Comparison on KO prediction task across 7 seeds of randomly splitting data into train-test splits. paired t-tests, *p <* 0.001). These results indicate that Nammu’s protein representations capture functional information relevant to KO classification in newly sequenced environmental metagenomes, even in the absence of surrounding genomic context.

## 4 Discussion and Limitations

We introduce Nammu, a Mamba-based foundation model for metagenomic sequences that integrates nucleotide and protein representations within extended genomic contexts. Building on advances in bidirectional state space models and mixed-modality genomic language modeling, Nammu is, to our knowledge, the first state space–based genomic language model designed for long-context mixed nucleotide–protein metagenomic representation learning. Across diverse downstream tasks, including taxonomic classification, functional annotation, biosynthetic gene cluster classification, and operon characterization, Nammu demonstrated the potential of state space architectures to learn biologically informative representations from metagenomic sequences. These findings also support the OMG corpus as a resource for large-scale genomic foundation model pretraining.

Nammu’s performance across these benchmarks highlights the potential value of genomic context for biological representation learning. In the CAMI II Marine dataset, Nammu generally outperformed gLM2 in taxonomic classification. However, this comparison evaluated held-out contigs from genomes that were closely related to the training set, rather than assessing generalization to phylogenetically distant or previously unseen organisms. Further evaluation on poorly characterized microbial lineages that are phylogenetically distant from those represented in the training data will be essential to determine Nammu’s ability to generalize beyond known microbial diversity.

Beyond taxonomic classification, we explored potential applications of Nammu’s representations across several biologically distinct tasks. For protein functional annotation in environmental deep-sea metagenomes, Nammu achieved consistent improvements over gLM2. More pronounced improvements were observed in operon-related tasks, particularly enzyme function prediction using operon-level context, where Nammu substantially outperformed gLM2. Importantly, these experiments were designed to evaluate the information captured by genomic foundation models rather than provide comprehensive benchmarking against established annotation tools such as BLAST. Such comparisons remain an important direction for future work. Interestingly, contextual genomic representations also enable tasks that are difficult to address through sequence alignment alone, such as identifying operon boundaries from intergenic regions and their surrounding coding sequences, potentially opening new avenues for downstream biological applications. Together, these results highlight the potential of mixed-modality genomic representations to support functional characterization beyond conventional sequence similarity-based approaches.

Our team recognizes several limitations of this study that should be considered when interpreting these findings. Our benchmarking primarily focused on comparisons with gLM2, without including established protein language models such as ESM-2. Although gLM2 provides a relevant baseline for contextual genomic representation learning, comparisons with protein-only foundation models would help establish whether mixed-modality representations provide additional information beyond protein sequence alone. Furthermore, Nammu and gLM2 differ in architecture, pretraining strategy, and context length, preventing us from attributing performance improvements to any individual component. While several evaluations incorporated taxonomic separation, further testing on previously unseen organisms and diverse environmental datasets will be important to better understand the scope and limitations of Nammu’s applicability. In addition, although the operon benchmarks demonstrate the predictive value of Nammu’s representations, further controlled comparisons between isolated proteins and proteins embedded within their genomic context are needed to quantify the specific contribution of surrounding sequences.

An important advantage of state space architectures is their potential computational efficiency when processing long sequences (Gu and Dao, 2024). Nammu was trained with a context window of 20,000 tokens, substantially exceeding the 4,096-token context of gLM2-150M and allowing larger genomic regions to be represented within a single input. Although previous studies suggest that Mamba-based architectures may generalize beyond their training context lengths (Gu and Dao, 2024; Wang et al., 2024), this capability has not yet been established for Nammu. Future work should systematically investigate context-length scaling, model size, and newer state space architectures (Dao and Gu, 2024; Lahoti et al., 2026). Additionally, our current training configuration did not explicitly prevent padding tokens from being processed by the state space layers. Incorporating sequence packing or state masking could improve training efficiency by reducing unnecessary state updates, providing an opportunity to further optimize Nammu’s architecture and performance.

More broadly, Nammu provides a foundation for extending contextual representation learning to additional challenges in microbiome research. Potential applications include functional characterization of poorly annotated proteins, identification of relationships across distant genomic regions, and refinement of metagenomic bins in complex microbial communities. Future approaches could also integrate representations across nonadjacent genomic regions or multiple contigs, moving beyond the contiguous sequence context currently modeled by Nammu. Together with recent advances in genomic language modeling, including gLM2, our findings motivate further development of long-context, mixed-modality architectures for investigating microbial diversity and function.

## Data Availability

The OpenMetaGenomic (OMG) corpus, CAMI II, MIBiG 4.0 and DGEB datasets used in this study are publicly available from their original sources cited in the Methods. Code is available at https://github.com/MaudeDavidLab/nammu. All processed contigs used for functional annotation are available at https://files.cqls.oregonstate.edu/David_Lab/nammu/.

## Author Contributions

N.A. developed and implemented Nammu, performed model training and benchmarking, analyzed the results, and contributed to manuscript preparation. J.H.W. contributed to data curation and model development. T.S. implemented and performed the operon classification benchmarks. K.I. conducted the large-scale taxonomic classification benchmark using the CAMI II Strain Madness dataset and Kraken2. A.R.T. contributed to biological interpretation and research supervision. M.D. conceived and supervised the study, secured funding, and contributed to study design, interpretation of results, and manuscript preparation. All authors reviewed and approved the final manuscript.

## Acknowledgments

We thank the Center for Quantitative Life Sciences (CQLS) at Oregon State University for their technical support and assistance with the NVIDIA GH200 computing infrastructure used in this study.

## Availability

Code is available at https://github.com/MaudeDavidLab/nammu and data at https://files.cqls.oregonstate.edu/David_Lab/nammu/.

