## Supplemental for "Metagenomic contextualization of proteins with state space models"

### Supporting information for:
### Nammu: Metagenomic Contextualization of Proteins Using State Space Models

Nima Azbijari[1,2*] ,Jacob H. Wynne[3] , Kristopher Irizarry[1,2] ,Tanush Sistla[4], Andrew R. Thurber[5,6] and Maude David[2]

[1]Department of Electrical Engineering and Computer Science, Oregon State University, Corvallis, OR, USA

[2]Department of Microbiology, Oregon State University, Corvallis, OR, USA

[3]Interdepartmental Graduate Program in Marine Science, University of California, Santa Barbara, Santa Barbara, CA, USA

[4]Department of Biochemistry and Biophysics, Oregon State University, Corvallis, OR, USA

[5]Marine Science Institute, University of California, Santa Barbara, Santa Barbara, CA, USA

[6]Department of Ecology, Evolution and Marine Biology, University of California, Santa Barbara, Santa Barbara, CA, USA

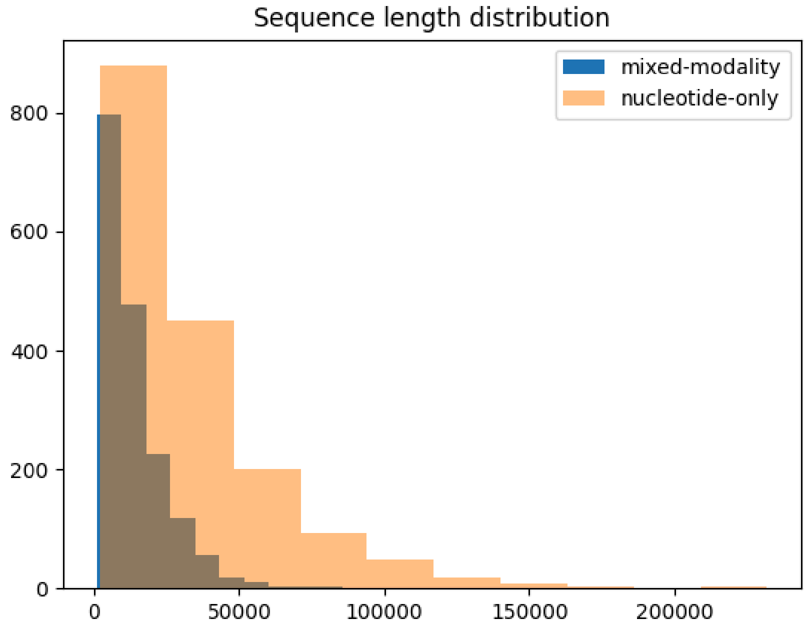


**Supplementary Figure 1 : Sequence length distributions of biosynthetic gene clusters (BGCs)**. Distribution of BGC sequence lengths for mixed-modality and nucleotide-only representations. BGCs span a wide range of sequence lengths, with nucleotide-only representations extending to substantially longer sequences, highlighting the importance of long-context modeling for BGC analysis.

**Supplementary Table 1: BGC class prediction from a MIBiG contig (n=1706, 6 classes), scored over pooled out-of-fold predictions of a 5-fold split grouped by taxon.** Each cell is the mean over 3 seeds with Δ = max − min across those seeds in parentheses. Layer: hidden layer the embedding is pooled from (final, a middle layer, or layer 9). Best mean per metric in bold. Superscripts on all metric columns mark a significant difference (paired t-test over the 3 seeds, which share folds across representations; Holm-corrected $p < 0.05$ over the 40 tests of each metric) from: a, the best k-mer row for that metric and classifier (†); b, the same model and layer on nt-only input; c, the same model and input at the final layer; d, the gLM2 row with matching input, layer type and classifier. Direction is read from the values.

| Representation | Input | Layer | $d$ | Clf. | Accuracy (Δ) | Macro-F1 (Δ) | MCC (Δ) | Macro-AUPRC (Δ) |
|---|---|---|---|---|---|---|---|---|
| Majority class | — | — | — | — | 0.312 | 0.079 | 0.000 | 0.167 |
| *Context-free baseline* | | | | | | | | |
| $k$-mer ($k$=4) | nt | — | 256 | kNN | 0.490$^{\dagger}$ (0.003) | 0.374$^{\dagger}$ (0.022) | 0.330$^{\dagger}$ (0.004) | 0.385$^{\dagger}$ (0.008) |
| $k$-mer ($k$=4) | nt | — | 256 | RF | 0.550 (0.015) | 0.418$^{\dagger}$ (0.016) | 0.411 (0.020) | 0.524$^{\dagger}$ (0.002) |
| $k$-mer ($k$=6) | nt | — | 4096 | kNN | 0.430 (0.007) | 0.276 (0.000) | 0.245 (0.009) | 0.347 (0.002) |
| $k$-mer ($k$=6) | nt | — | 4096 | RF | 0.562 (0.007) | 0.405 (0.009) | 0.426 (0.010) | 0.518 (0.014) |
| $k$-mer ($k$=8) | nt | — | 65536 | kNN | 0.373 (0.009) | 0.152 (0.004) | 0.163 (0.020) | 0.271 (0.007) |
| $k$-mer ($k$=8) | nt | — | 65536 | RF | 0.577$^{\dagger}$ (0.009) | 0.398 (0.021) | 0.447$^{\dagger}$ (0.013) | 0.511 (0.009) |
| *gLM2 (150M)* | | | | | | | | |
| gLM2-150M | nt only | final | 640 | kNN | 0.458$^{a}$ (0.003) | 0.362 (0.010) | 0.285$^{a}$ (0.005) | 0.341 (0.015) |
| gLM2-150M | nt only | final | 640 | RF | 0.530 (0.009) | 0.436 (0.012) | 0.383 (0.013) | 0.489 (0.015) |
| gLM2-150M | mixed | final | 640 | kNN | 0.626$^{a,b}$ (0.004) | 0.512$^{b}$ (0.004) | 0.515$^{a,b}$ (0.005) | 0.528$^{a,b}$ (0.007) |
| gLM2-150M | mixed | final | 640 | RF | 0.691$^{a,b}$ (0.005) | 0.588$^{a,b}$ (0.004) | 0.600$^{a,b}$ (0.006) | 0.674$^{a,b}$ (0.007) |
| gLM2-150M | nt only | mid | 640 | kNN | 0.464 (0.005) | 0.348 (0.014) | 0.294 (0.006) | 0.337 (0.005) |
| gLM2-150M | nt only | mid | 640 | RF | 0.534 (0.008) | 0.417 (0.019) | 0.388 (0.011) | 0.468$^{a}$ (0.007) |
| gLM2-150M | mixed | mid | 640 | kNN | 0.763$^{a,b,c}$ (0.014) | 0.692$^{a,b,c}$ (0.020) | 0.697$^{a,b,c}$ (0.019) | 0.727$^{a,b,c}$ (0.015) |
| gLM2-150M | mixed | mid | 640 | RF | **0.834**$^{a,b,c}$ (0.004) | 0.769$^{a,b,c}$ (0.009) | **0.789**$^{a,b,c}$ (0.005) | 0.828$^{a,b,c}$ (0.016) |
| *NAMMU* | | | | | | | | |
| NAMMU | nt only | final | 1024 | kNN | 0.458 (0.009) | 0.353 (0.014) | 0.286 (0.013) | 0.337 (0.009) |
| NAMMU | nt only | final | 1024 | RF | 0.519 (0.011) | 0.387$^{d}$ (0.012) | 0.367 (0.015) | 0.444$^{a}$ (0.002) |
| NAMMU | mixed | final | 1024 | kNN | 0.659$^{a,b}$ (0.015) | 0.541$^{b}$ (0.020) | 0.561$^{a,b}$ (0.022) | 0.562$^{a,b}$ (0.017) |
| NAMMU | mixed | final | 1024 | RF | 0.729$^{a,b}$ (0.008) | 0.632$^{a,b}$ (0.011) | 0.650$^{a,b}$ (0.011) | 0.719$^{a,b,d}$ (0.005) |
| NAMMU | nt only | 9 | 1024 | kNN | 0.454 (0.010) | 0.351 (0.013) | 0.282 (0.014) | 0.336 (0.006) |
| NAMMU | nt only | 9 | 1024 | RF | 0.528 (0.013) | 0.400 (0.017) | 0.381 (0.018) | 0.451$^{a}$ (0.008) |
| NAMMU | mixed | 9 | 1024 | kNN | 0.776$^{a,b,c}$ (0.010) | 0.730$^{a,b,c}$ (0.015) | 0.717$^{a,b,c}$ (0.013) | 0.763$^{a,b,c}$ (0.006) |
| NAMMU | mixed | 9 | 1024 | RF | 0.826$^{a,b,c}$ (0.004) | **0.781**$^{a,b,c}$ (0.007) | 0.777$^{a,b,c}$ (0.005) | **0.860**$^{a,b,c}$ (0.002) |
